# A computational method to aid the design and analysis of single cell RNA-seq experiments for cell type identification

**DOI:** 10.1101/247114

**Authors:** Douglas Abrams, Parveen Kumar, R. Krishna Murthy Karuturi, Joshy George

**Affiliations:** Colby College, Waterville, Maine, USA.; The Jackson Laboratory for Genomic Medicine, Farmington, CT, USA 06030.

**Keywords:** single cell RNA-seq, cell-type identification, clustering

## Abstract

**Background:** The advent of single cell RNA sequencing (scRNA-seq) enabled researchers to study transcriptomic activity within individual cells and identify inherent cell types in the sample. Although numerous computational tools have been developed to analyze single cell transcriptomes, there are no published studies and analytical packages available to guide experimental design and to devise suitable analysis procedure for cell type identification.

**Results:** We have developed an empirical methodology to address this important gap in single cell experimental design and analysis into an easy-to-use tool called SCEED (Single Cell Empirical Experimental Design and analysis). With SCEED, user can choose a variety of combinations of tools for analysis, conduct performance analysis of analytical procedures and choose the best procedure, and estimate sample size (number of cells to be profiled) required for a given analytical procedure at varying levels of cell type rarity and other experimental parameters. Using SCEED, we examined 3 single cell algorithms using 48 simulated single cell datasets that were generated for varying number of cell types and their proportions, number of genes expressed per cell, number of marker genes and their fold change, and number of single cells successfully profiled in the experiment.

**Conclusions:** Based on our study, we found that when marker genes are expressed at fold change of 4 or more than the rest of the genes, either Seurat or Simlr algorithm can be used to analyze single cell dataset for any number of single cells isolated (minimum 1000 single cells were tested). However, when marker genes are expected to be only up to fC 2 upregulated, choice of the single cell algorithm is dependent on the number of single cells isolated and proportion of rare cell type to be identified. In conclusion, our work allows the assessment of various single cell methods and also aids in examining the single cell experimental design.

## Background

The greater precision afforded by single cell sequencing has increased the scope of the average sequencing study. Unlike conventional methods that profile hundreds or thousands of cells (aka bulk sequencing or profiling), the advancement in next generation sequencing and microfluidic technologies have now made it possible to isolate a single cell and perform different types of omics profiling including genomics, transcriptomics, epigenomics and proteomics [1]. One prominent technique that measures gene expression at single-cell level is single cell mRNA sequencing (scRNA-seq) [1, 2]. It, unlike bulk sequencing, unmasks the fundamental, widespread heterogeneity in gene expression among cells in a tissue or cells considered to be of same type based on canonical markers [3, 4]. Hence, rather than simply examining differential expression between two samples, we can identify the cell types and expressed genes within each cell type as a first step before differential expression analysis [4, 5]. Not only does this first step provide valuable insights into the transcriptomic profiles of individual cell types and states, but it also provides a deeper context for the subsequent differential expression analysis.

However, the effectiveness of cell type identification is a multi-step process which led to the explosion of new single cell software applications, referred to as a “cottage industry” [6]. According to *Awesome Single Cell* (https://github.com/seandavi/awesome-single-cell), a site that compiles a list of new single cell analysis methods, 89 methods have recently been created for analyzing single cell sequencing data (normalization, dimensionality reduction, clustering and differential expression), including plethora of methods required of cell type identification.

Hence, it is necessary to comparatively assess the different tool combinations (aka pipelines) to determine which is best at cell type identification. Comparative analyses have been published on sequencing [7, 8], normalization [9] and clustering [10, 11]. Yet, there has not been a comprehensive study, assessing whole pipelines and addressing broader issues of experimental design in cell type identification.

We developed a computational method to address this important gap. We developed an easy to use tool in an R-package SCEED (Single Cell Experimental Design and Analysis). The package has functionality to simulate scRNA-seq data with user provided statistical characteristics: total number of cells, genes, groups proportions, marker genes and fold change (fC) of marker genes. The simulated dataset with known cell types can be analyzed using published cell-type identification algorithms using SCEED. Systematic comparison of the results of the analysis pipeline to the known true labels using accuracy, sensitivity, specificity, F1score and Rand Index (for details see methods) that provide the ability to identify the optimal single cell algorithms for dataset and will also help to identify the number of cells required for adequate power for the detection of the cell-types.

## Methods

The schematic of SCEED is shown in Figure 1. Each step in SCEED is described below.

**Figure 1.**
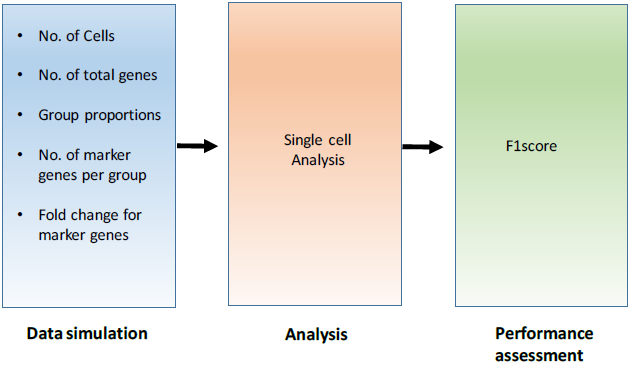
Schematic representation of SCEED pipeline. (Left to right) First a simulated dataset is generated using SCEED “generateDataset” function with input parameters mentioned under “Data simulation”. Next, the simulated dataset is analyzed using different single cell algorithms. To test the performance of each single-cell algorithm, F1-score which is a measure of test’s accuracy is computed. Finally, based on the F1 score cutoff chosen by user, best algorithm and number of cells required to perform the single cell experiment are selected.

## Data simulation

Our procedure to simulate the single cell data is shown in Figure 2. In step 1, gene by cell expression matrix is simulated using *Splatter* package [12], which simulates *m* cell types of given rarity/prevalence with *n* cells. In step 2, each cell type will express specific number of marker genes *g* with specific fold change levels *f*_*c*_. The mean expression level of each marker gene *g*_*i*_ in group *k* was simulated by taking the product of a group-specific fold change level (sampled from a negative binomial distribution with shape= *f*_*ci*_ and rate= 1) and the mean expression level of *g*_*i*_ in all cells that are not part of *k*. For each cell in *k*, the final expression level of marker gene *g*_*i*_ was the product of the simulated mean of *g*_*i*_ and a library size that was simulated using Splatter [12]. The remaining steps are stated in Figure 2.

**Figure 2:**
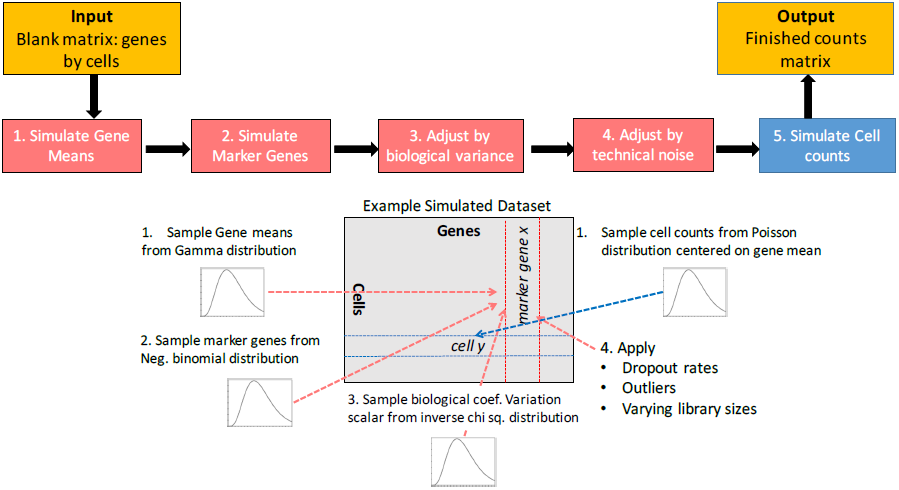
Generation of simulated dataset

## Analyses

### 1. Single cell Analysis Steps

A standard single cell analysis procedure includes data normalization, dimensionality reduction and clustering [13]. Normalization is a crucial step for any single cell analysis that adjusts for unwanted technical or biological variations that may affect the gene expression analysis. With larger datasets like single cell, dimensionality reduction is also an important step that transforms data into lower dimensional sub-space, allowing significant reduction in data complexity and also makes data visualization easier. Finally, single cells with similar transcriptome profiles are clustered together to deduce putative (sub)populations, aka cell types.

### 2. Incorporation of single cell methods into SCEED package

SCEED package allow users to add any single cell analysis package of interest into its pipeline using function “sceed_*AlgorithmName*” for example sceed_seurat. For example, in the current implementation of SCEED, *kmeans*, *Simlr* and *Seurat* (details in results section) are available. Although we have added only three single cell algorithms, SCEED package is completely flexible and any number of single-cell algorithms can be added for testing as per user’s requirements.

## Performance assessment

The performance of an analysis procedure is assessed by computing F1 score of a cluster. F1score is a balancing measures of recall (sensitivity) and precision of cell classification. Higher F1 score shows better performance of the algorithm tested. User can choose F1-score threshold suitable to annotate the clusters for cell types and hence best single cell analysis algorithm as well as sample size.

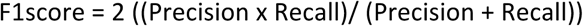

## Results

We used SCEED to test 3 popularly known single cell algorithms for cell type identification: k-means, SEURAT and SIMLR. For k-means clustering approach, k was set equal to number of cell types simulated. For Seurat and Simlr algorithms, default parameters mentioned by the authors were used. In Seurat, while using “FindClsuters” function, k.param was set to the number of cell types simulated. We generated 27 datasets of varying choices of parameters.

### Generating simulated single-cell datasets

In a single cell experiment, discovering rare cell population is of utmost importance. Stressing on the rarity of cell populations, we simulated single cell datasets where five cell types were partitioned into unequal proportions such that one of them has low proportion or representing rare population, ranging from 2%-10%. For instance, we defined a single-cell category having 5 cell types in proportions of 0.1, 0.2, 0.2, 0.2 and 0.3. In each cell type, 50 genes were simulated as marker genes that were either 2-, 4- or 8-fold upregulated when compared to rest of the cell types. For the same proportions of cell types while keeping the other parameters same, we simulated single cell data sets of 2000 or 3000 cells. More details of each dataset are shown in Table 1. In summary, we created 27 simulated single-cell datasets. Notably, in SCEED package, all these parameters (such as number of cell types, single cells per cell type, genes per cell, marker genes per cell type and fold change cutoffs) can be adjusted as per user’s requirements.

**Table 1:** Properties of different of simulated single cell datasets generated.

| Cell type proportions | No. of cell types (m) | No. of Genes | No. of Marker genes | No. of cells simulated (n) | Fold change (fC) of marker genes |
| --- | --- | --- | --- | --- | --- |
| 0.1, 0.2, 0.2, 0.2, 0.3 | 5 | 10000 | 50 | 1000, 2000 and 3000 | 2, 4 and 8 |
| 0.05, 0.2, 0.2, 0.2, 0.35 | 5 | 10000 | 50 | 1000, 2000 and 3000 | 2, 4 and 8 |
| 0.02, 0.2, 0.2, 0.2, 0.38 | 5 | 10000 | 50 | 1000, 2000 and 3000 | 2, 4 and 8 |

### Testing the performance of single-cell algorithms and estimation of sample size required

All these datasets were analyzed using three single cell algorithms, kmeans, Seurat and Simlr and tested for their performance using F1 score. At lowest fold change (fC) of 2 for marker genes, irrespective of number of single cells collected, Seurat provided the best performance in F1-score for rarity of 0.1. However, for fc of 2, we may need at least 1000 cells for F1 score of >0.9. As fC increases, the other algorithms also offered increased performance, Supp figures 1 and 2. Next, we compared these algorithms to detect even rarer cell type, with a proportion of 0.05 (the cell type proportions are 0.05, 0.35, 0.2, 0.2 and 0.2), fig 3. At fC 2, Seurat reached the F1 score of 0.93 but only when number of single cells >= 2000, Fig 3. In line with previous observation, the other algorithms also showed increased performance with increase in fC at 0.05 proportion, Supp fig 1,2. However, when we reduced the rarer cell type proportion further down to 0.02, Simlr outperformed the remaining two algorithms with F1 score of 0.69 for number of single cells >= 1000. Separately, we also estimated the minimum sample size required at a given F1 score. For instance, Simlr could reach to F1 score of >0.7 for proportions of 0.1 and 0.05 for sample size (number of single cells) of 1000 while Seurat required sample sizes of 1000 and 2000 for cell proportions of 0.1 and 0.05 respectively, Supp fig 3.

**Figure 3:**
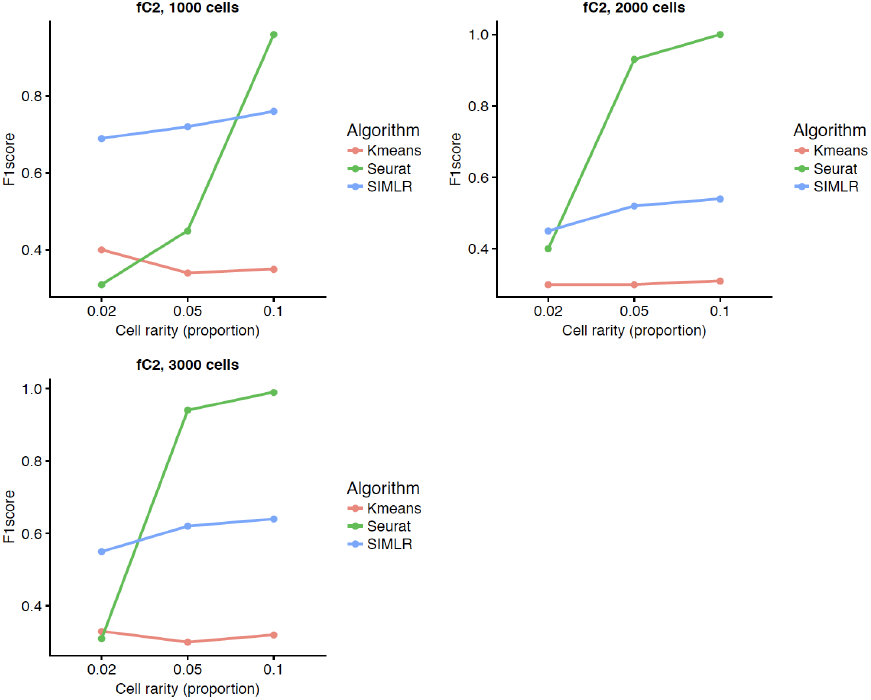
Performance of different single cell algorithms at different cell proportions.

## Discussion and Conclusion

We proposed SCEED method as an easy-to-use package to help the researchers in designing a single cell experiment (estimate the number of cells required to identify novel cell types) and optimal analysis procedure. The package takes into account all technical and biological parameter that characterize typical single cell RNA-seq data. Using SCEED package, we simulated 27 single cell datasets that account for varying sample sizes, rarity of cell types and fold change of expression of marker genes. Such a simulation is significant. For example, when researchers are planning to analyze cell types similar to beta cells from islets of Langerhans in the pancreas where marker genes such as insulin are expressed in far greater concentrations than rest of the genes. In contrast, they are interested in identifying sub classes of established cell types like dendritic cells. Using SCEED package, researchers can generate simulated datasets that bear statistical properties similar to that of the expected data and test various single cell algorithms. Our package not only suggest the best method among the tested algorithms but also suggest the number of cells required to achieve the required results. As single cell transcriptome analysis field is rapidly growing field, SCEED package facilitates easily adding more single cell algorithms for testing.

In our study, we have compared the performance of three popularly used single cell algorithms. Though our simulations are limited, our study clearly shown that even popularly used algorithms do not perform best over ranges of cell population rarity and fold change in expression of marker genes. Based on these results, we demonstrated that SCEED package fills an important gap in the single cell analysis field. However, we need to conduct extensive study to identify optimal analysis procedures for a variety of experimental settings and statistical properties of data. Such a study needs to account not only for the 3 parameters we tested up on, it needs to account for the variation in the other statistical parameters (can be selected in SCEED package) and addressing the experimental designs of scRNA-seq experiments.

## Authors contributions

J.G. and K.K. conceptualized the project and methodology. D.A., P.K. and J.G. analyzed the data. D.A., P.K., J.G. and K.K. wrote the manuscript. D.A. and P.K. created the figures. All authors read and approved the manuscript.

## Acknowledgements

We thank Kyung In Kim for his valuable inputs for the manuscript. We thank National Cancer Institute of the National Institutes of Health for Summer Student Fellowship under award number R25 CA174584 to Douglas Abrams.

